# Leaf trichome diversity, acylsugar concentration, and their relationships to leaf area in *Solanum galapagense*

**DOI:** 10.1101/2022.07.19.500675

**Authors:** Ilan Henzler, Hamid Khazaei

## Abstract

Glandular trichomes are physical and chemical barriers used by some tomato wild relatives to confer resistance against insect pests and diseases transmitted by them. *Solanum galapagense* has been identified as one of the potential sources of insect pest resistance. The present study aimed to examine the trichome diversity and acylsugar concentration of 26 accessions of *S. galapagense* along with one cultivated tomato (*S. lycopersicum*) and one cherry tomato (*S. l. cerasiforme*) cultivar. The results revealed large genetic variation among *S. galapagense* accessions for all studied traits. The *S. galapagense* accessions had significantly higher trichome types IV on the adaxial and abaxial surfaces of the leaf and greater acylsugar concentration but smaller leaflet area than cultivated tomato. The selected cherry tomato line represents greater trichome type IV and acylsugar than other groups. The acylsugar concentration was positively associated with trichome type IV but negatively associated with trichome type V on both leaf surfaces. Leaflet area was negatively associated with trichome IV density and acylsugar concentration. Analysis of DNA markers revealed the presence of two previously identified whitefly-resistance alleles in *S. galapagense* accessions. This study will support breeding programs aiming to improve insect pest resistance in tomato cultivars using crop wild relatives.

## Introduction

Cultivated tomato (*Solanum lycopersicum*) is the most valuable vegetable crop by fruit weight globally, generating revenues of $70 billion from 187 million tonnes of fresh fruit in 2020 (FAOSTAT, 2022). Improving fruit quality and yield through domestication in this crop, has led to the loss of important plant defence characteristics (Paudel *et al*., 2019), with tomato cultivation now heavily relying on pesticides to counter biotic stress resistance (Dari *et al*., 2016). The chemical treatments are not only costly, but they also harm the environment (Damalas and Eleftherohorinos, 2011).

Developing resistant tomato cultivars could reduce the reliance on pesticides and their associated burden.

Tomato wild relatives are important sources of genetic diversity and are commonly used as reliable sources of resistance genes against abiotic and biotic stress (Ebert and Schafleitner, 2015; Khazaei and Madduri, 2022). Sources of resistance to insect pests have been identified in some tomato wild species including *S. galapagense, S. habrochaites, S. pennellii, S. cheesmaniae*, and *S. pimpinellifolium* (e.g. Kennedy 2003; Schilmiller *et al*., 2012; Leckie *et al*., 2016; Rakha *et al*., 2017b; Vosman *et al*., 2018). Among them, *S. galapagense* has been identified as one of the most promising sources of insect pest resistance (Firdaus *et al*., 2012; Lucatti *et al*., 2013). It has been the focus of most tomato breeding programs aiming to improve biotic and abiotic stress resistance due to its close relationship to cultivated tomato (Vendemiatti *et al*., 2021). The *S. galapagense* species originates from the Galápagos Islands, an archipelago 1000 km west of Ecuador where it formed a diverse range of phenotypes due to the islands’ unique ecosystem (Darwin *et al*., 2003). While genetic studies revealed a narrow genetic diversity within the *S. galapagense* germplasm (Darwin, 2009; Pailles *et al*., 2017), it presents distinct morphological characteristics. These include yellow-green foliage, orange fruit at maturity, small seed size, and highly divided leaves (Darwin *et al*., 2003; Fenstemaker *et al*., 2022).

Plants have developed a variety of defence mechanisms to counter biotic and abiotic stress conditions (e.g. Levin, 1973;Oksanen, 2018). One of these is the presence of hair-like structures, called trichomes, on the surface of flowers, fruits, stems, and leaves as physical and chemical lines of defence. Numerous studies have been conducted on the nature of these epidermal outgrowths including function, quantification, and effectiveness (*see* Glas *et al*., 2012; Vendemiatti *et al*., 2021). In tomato, among the seven types of trichomes found on plants, four are termed glandular due to the sugar-filled pouches at their tip (Luckwill, 1943). The presence of glandular trichome types IV and VI has been associated with higher insect pest resistance (Lucatti *et al*., 2013; Firdaus *et al*., 2013; Rakha *et al*., 2017b; Zhang *et al*., 2020). These types of glandular trichomes deter insects through the release of acylsugars which cause behaviour changes (antixenosis) and reduced survival (antibiosis) in the arthropods that land on them (Antonious *et al*., 2005; Bleeker *et al*., 2011, 2012; Dias *et al*.,2016).

Trichome diversity, density, and its relationship with insect pest resistance have been investigated in tomato wild relatives including *S. galapagense* (e.g. Firdaus *et al*., 2012, Lucatti *et al*., 2013; Rakha *et al*., 2017b). An aspect deserving further attention is to harness the genetic diversity of morphological and biochemical characteristics of large germplasm of *S. galapagense* accessions and their relationships with leaf area. So, this study aims to uncover differences in trichome diversity, leaf characteristics, and acylsugar concentration in this species. This is supported by the analysis of DNA markers associated with insect pest resistance phenotypes (Firdaus *et al*., 2013).

## Materials & Methods

### Plant material

This study was conducted on 27 accessions of *S. galapagense*, one accession of cherry tomato (*S. lycopersicum* var. *cerasiforme*, abbreviated as *S. l. cerasiforme)*, and one cultivated tomato (*S. lycopersicum)*. Detailed information on accession number, origin, and habitat at collection sites are presented in Table 1. All *S. galapagense* accessions originated from the Galápagos Islands, Ecuador (Figure 1). More than a third were collected on Isla Isabela, the largest island. Each accession’s habitat was unique which might point to the reason for the high degree of variation observed between them. The cultivated tomato is a breeding line from the World Vegetable Center (WorldVeg) carrying multiple tomato yellow leaf curl virus resistance genes (*Ty-1/3* and *Ty-2*). The SM131 cherry tomato is a selection from an accession from Galápagos Islands due to its high density of trichome type IV (unpublished data). All accessions were acquired from the WorldVeg genebank. Accession VI037324 did not germinate and was removed from experiments.

**Table 1.** Species, accession number, origin, and habitat at collection sites of *Solanum* species used in this study.

| Species | Accession No. | Origin | Elevation (m) | Other name(s) | Habitat/phenotype |
| --- | --- | --- | --- | --- | --- |
| *S. galapagense* | VI007099 | Bartolome, Galápagos Islands, Ecuador | 15 | LA0317, PI231257 | Lava flow, amongst basalt rock, very arid, coastal arid zone |
|  | VI037239 | Isabela, Galápagos Islands, Ecuador | 40 | LA0436 | Sandy, near lava outcrop |
|  | VI037241 | Pinta, Galápagos Islands, Ecuador | 150 | LA0526, SAL254 | West side Abingdon island |
|  | VI037324 | Isabela, Galápagos Islands, Ecuador | 50 | LA1410, PI365898 | Quebrada near landing |
|  | VI037339 | Isabela, Galápagos Islands, Ecuador | 5 | LA1401, PI365897 | Among rocks in large beach with magma and lava flows at each end, collected few metres above tide line |
|  | VI037340 | Isabela, Galápagos Islands, Ecuador | 200 | LA1408, PI379039 | Ridge of Cape Berkeley volcano |
|  | VI037867 | Floreana, Galápagos Islands, Ecuador | - | LA1136, TL01054 | Garnder near Floreana islet |
|  | VI037868 | Rabida, Galápagos Islands, Ecuador | 10 | LA1137, TL01055 | On cinder ash |
|  | VI037869 | Santiago, Galápagos Islands, Ecuador | 650 | LA1141, TL01056 | Interior walls of crater, purple fruit colour (Fenstermaker et al., 2022) |
|  | VI045262 | Santiago, Galápagos Islands, Ecuador | - | Selection from LA1141, TL01572 | - |
|  | VI057400 | Fernandina, Galápagos Islands, Ecuador | - | LA483, 6201A, SAL241 | - |
|  | VI057408 | Isabela, Galápagos Islands, Ecuador | - | Selection from LA1401 | - |
|  | VI057457 | Galápagos Islands, Ecuador | - | LA3909 | - |
|  | VI063173 | Bartolome, Galápagos Islands, Ecuador | 15 | LA0317 | Lava flow, amongst basalt rock, very arid |
|  | VI063174 | Isabela, Galápagos Islands, Ecuador | 30 | LA0438, SAL192 | Rocky basalt outcropping in first hills to W. (7 km) from Villamil, 1 km from coast |
|  | VI063175 | Isabela, Galápagos Islands, Ecuador | 20 | LA0480A, SAL238 | Along coast in bay facing Cowley Islet - not far from shore, in broken terrain without shade |
|  | VI063176 | Santa Cruz, Galápagos Islands, Ecuador | - | LA0528, SAL256 | Academy Bay |
|  | VI063177 | Fernandina, Galápagos Islands, Ecuador | 580 | LA0530, SAL258 | Inside crater at edge |
|  | VI063178 | Galápagos Islands, Ecuador | 5 | - | - |
|  | VI063179 | Santiago, Galápagos Islands, Ecuador | 6 | LA0747 | In lava formation near shore |
|  | VI063180 | Santiago, Galápagos Islands, Ecuador | 3 | LA0748 | In lava formation |
|  | VI063181 | Isabela, Galápagos Islands, Ecuador | 4 | LA0929 | Growing in lava, roots in sand, above sea level |
|  | VI063182 | Isabela, Galápagos Islands, Ecuador | 4 | LA0930 | Growing in lava, roots in sand, same as LA 929 |
|  | VI063183 | Bartolome, Galápagos Islands, Ecuador | - | LA1044 | - |
|  | VI063184 | Isabela, Galápagos Islands, Ecuador | 400 | LA1452 | On the trail from Punta Ecuador to crater rim - mid-elevation in longer of two lava flows |
|  | VI063185 | Isabela, Galápagos Islands, Ecuador | 100 | LA1627 | Volcanic cone above Darwin's salt lake - likely Tagus Cove |
|  | VI063187* | Santiago, Galápagos Islands, Ecuador | 10 | LA1411, PI379040 | On soft bright red rock formation, margin of beach |
| *S. l. cerasiforme* | SM131 | Fernandina, Galápagos Islands, Ecuador | - | - | - |
| *S. lycopersicum* | CLN3682C | Breeding line | - | AVT01424 | Pedigree: CLN3682F1-10-3-4-27-3-16 |

More information about accessions can be found at https://genebank.worldveg.org/ (accessions having "VI" code) and http://www.ars-grin.gov/ (accessions having "PI" code).
Passport data was obtained for accessions with "LA" code from the TGRC, http://tgrc.ucdavis.edu/.
\* This accession first was classified as *S. cheesmaniae* and later reclassified as *S. galapagense*.

**Figure 1.**
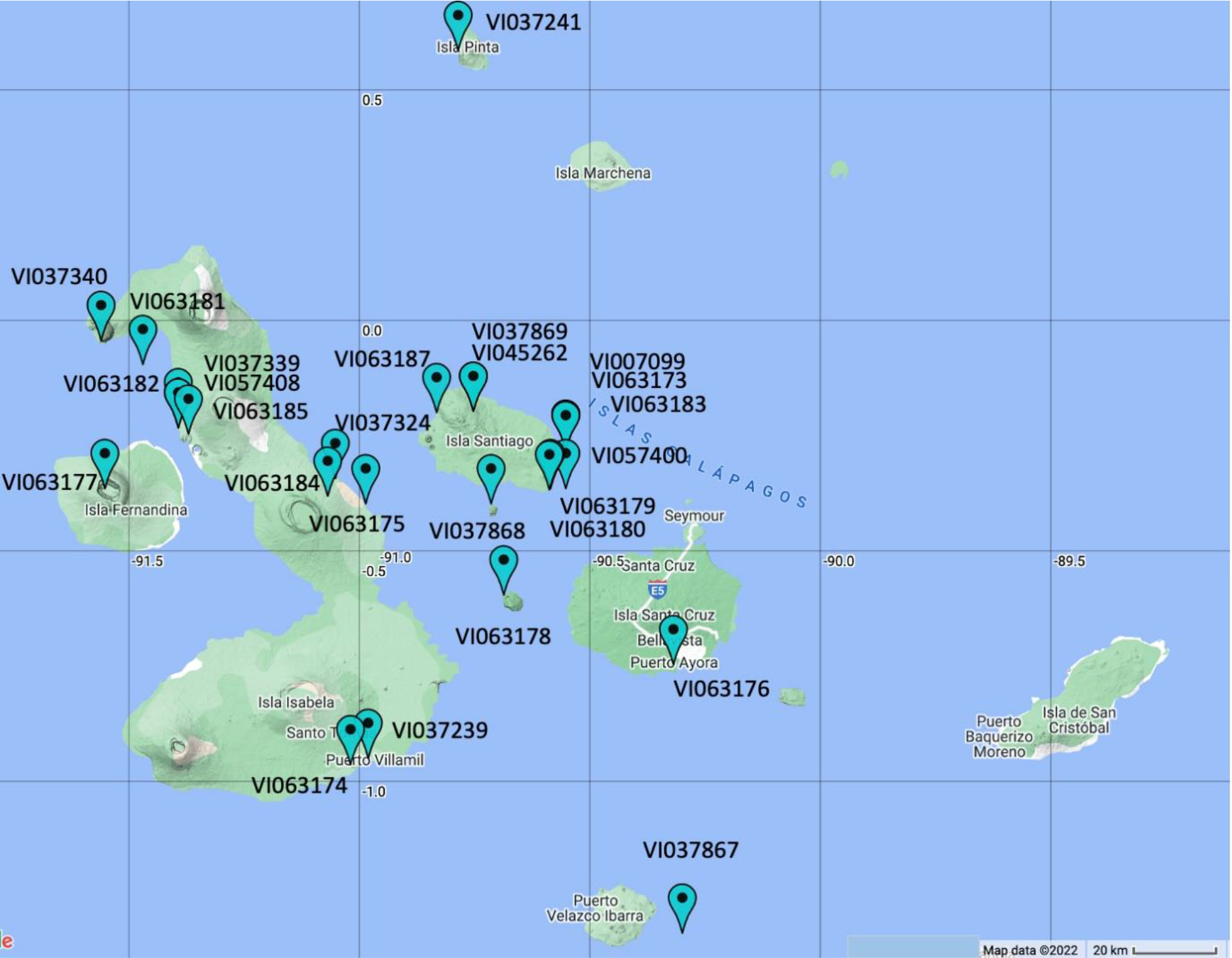
Geographical distribution of *S. galapagense* accessions in the Galápagos Islands.

### Seed treatment

Tomato seeds acquired from the WorldVeg genebank were treated with Hydrogen Chloride for 15 minutes and washed under running water. They were then treated with Trisodium Phosphate for 1 hour, washed under running water, and dried in an incubator room at 60% for two days.

### Growing conditions

Experiments were conducted in a glasshouse at the WorldVeg in Shanhua, Taiwan. Seeds were sown in a nursery on the 25th of February 2022, and after two weeks were transplanted into 8-inch pots filled with cultivable soil collected from tomato fields. The pots were arranged in a randomised complete block design with four replicates. Plants were watered once a day in the morning and fertilised with a blend of 15–15–15 (N–P–K) at week 4 after transplanting. Relative humidity was about 80±15%. The temperature was set to 28±3°C during the day and to 22±2°C during the night.

### Measurements

#### Leaflet area

The leaflet area was measured 10 weeks after sowing using the third leaf from the apex. It was measured using a LI-3100 leaf area meter (LI-COR Inc., Lincoln, NE, USA). The same leaflet was also used for subsequent trichome and acylsugar measurements.

#### Trichome Analysis

Analysis of leaf trichomes was conducted eight and 10 weeks after sowing using a stereo microscope (Leica^®^ M-Series Stereo microscopes, Ernst Leitz Wetzlar, GmbH, Germany). Leaf samples were collected at the third node from the apex using sterile forceps. The density of glandular trichome types I, IV, V, and VI were measured from four randomly chosen leaflets within 1mm^2^. The number of trichomes was counted from four different microscopic fields at 5X magnification and converted to the number per mm^2^ using a standard scale. The identification of trichome types on the leaf surface followed a schematic drawn by Luckwill (1943). After measuring trichomes on the adaxial surface, the leaflets were flipped, to measure trichomes on the abaxial surface.

#### Acylsugar concentration

Analysis of acylsugar content was conducted at eight and 10 weeks after sowing. Polyethylene vials were used to collect four 3±1 cm lateral leaflets from each plant at the third node from the apex. Samples were dried in an incubator at 29°C for 3 days before washing them with 3 ml methanol. Of this suspension, 100 μl was added to 100μl 6M Ammonium Hydroxide in 96 well ELISA plates with two biological replicates, following a protocol developed by Martha Mutschler (Savory, 2004). The samples were incubated overnight and left to dry under the hood for 3 days before adding 200 μl PGO reagent to each well and placing it on an orbital shaker. After 3 hours, absorbance values at 490 nm were measured using BioTek’s uQuant (Agilent Technologies Inc., Santa Clara, CA, USA) and converted into acylsugar concentration using a sucrose standard curve.

### DNA extraction and DNA marker assay

Genomic DNA was extracted from 10-week-old plants using the CTAB method (Doyle and Doyle, 1990). The *Wf-1* and *Wf-2* QTLs sequences presented in Firdaus *et al*. (2013) were used to genotype the studied germplasm for the presence/absence of corresponding bands. Purified DNA samples were digested with restriction enzymes DDeI and HpyCH4IV for *Wf-1* and *Wf-2* markers, respectively. Digested samples were amplified along with marker-specific primers using PCR reactions as described by Mahfouze and Mahfouze (2019). The PCR amplified samples were run on a 5% acrylamide gel for 30 minutes at 100V and stained using an ETBR-out stain. The gel was scanned in a Bio-1000F scanner, and the amplified bands visualised using Microtek MiBio Fluo software (both from MicroTek International, Inc., Hsinchu, Taiwan).

### Statistical analysis

The R statistical package (R Development Core Team, 2021) was used for data analysis. Correlation analysis was performed to determine the relationships between morphological measurements. The dataset was subjected to a one-way analysis of variance (ANOVA) and the SEM (standard error of means) was calculated. Principal component analysis (PCA) was employed to illustrate relationships between accessions and leaf morphological measurements. The online mapping tool at maps.co was used to plot the coordinates of accessions in Figure 1 (https://maps.co/). The GPS coordinates of *S. galapagense* accessions were obtained from the Tomato Genetics Resource Center (C.M. Rick TGRC, https://tgrc.ucdavis.edu/) and the WorldVeg (https://genebank.worldveg.org/#/) genebank databases.

## Results

Trichome densities varied significantly among studied germplasm (*P*<0.001). In *S. galapagense* accessions, trichome type I ranged from 0.0 to 1.5 for the abaxial and from 0.0 to 4.5 for the adaxial surface. Overall, *S. galapagense* accessions had significantly higher density of trichome type I (40% on abaxial and 550% on adaxial surfaces) than cultivated tomato (CLN3682C). The cherry tomato (SM131) had 157% more trichome type I on the abaxial and 10% less on the adaxial surface than the mean *S. galapagense* (Table 2). Trichome type IV density ranged from 6.3 to 13.5 for abaxial and 0.7 to 10.9 for the adaxial surface of *S. galapagense* accessions, while the cultivated tomato had none on either surface. The cherry tomato had 22% greater (on both surfaces) trichome type IV compared to the average values of *S. galapagense* accessions. The cultivated tomato had high levels of trichome type V with an average count of 17 on the abaxial and 9.5 on the adaxial surface while *S. galapagense* had few ranging from 0.0 to 6.1 on the abaxial and 0.0 to 9.0 on the adaxial surface (Table 2). The cherry tomato had higher counts of trichome type V than *S. galapagense* accessions with an increase of 520% on the abaxial and 416% on the adaxial surface. Within *S. galapagense* accessions, trichome type VI varied from 0.4 to 2.7 on the abaxial and 0.3 to 2.8 on the adaxial side. For the abaxial surface, this was 30% and 14% lower than cultivated and cherry tomato cultivars, respectively. For the adaxial surface, it was 94% lower and 30% higher than cultivated and cherry tomatoes, respectively (Table 2). As a general trend, most observed accessions showed fewer trichomes on the adaxial than on the abaxial side with 28% less for trichome type I, 13% less for type IV, 10% less for type V, and 40% less for type VI. Comparing 8- and 10-week trichome phenotyping at the abaxial surface, *S. galapagense* trichome densities remained stable with a 3% decrease for type I, 3% increase for type IV, and 12% decrease in type VI.

**Table 2.**
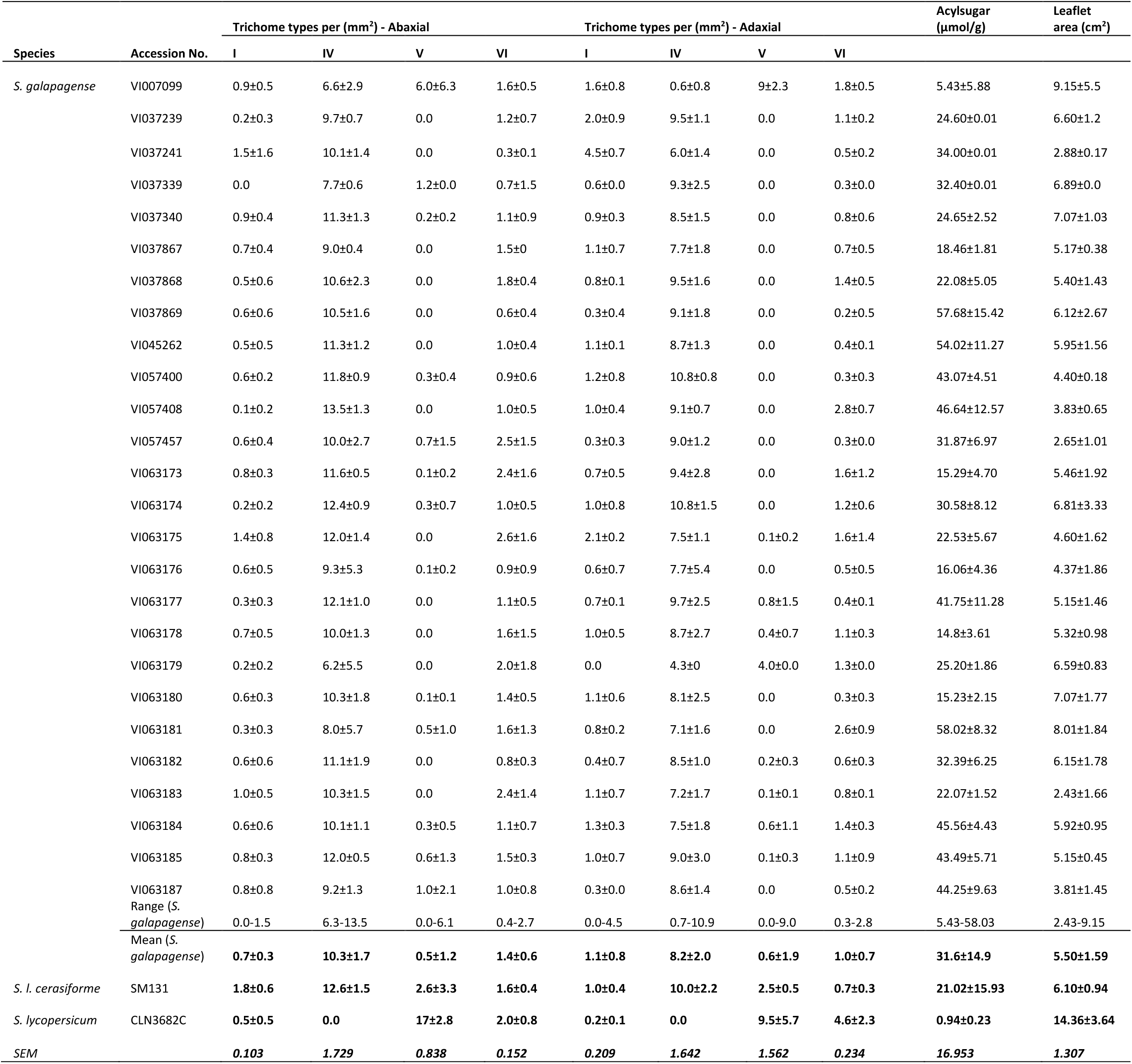
Mean ± SD (standard deviation) for trichome measurements (abaxial and adaxial surfaces) at 10 weeks after sowing and acylsugar concentration and leaflet area on 26 *S. galapagense* accessions along with one cherry tomato and one cultivated tomato genotype. SEM, standard error of means.

Acylsugar concentration varied significantly (*P*<0.001) between *S. galapagense* accessions, ranging from 5.43 to 58.03 μmol/g. The cultivated tomato cultivar (CLN3682C) showed a very low acylsugar concentration of 0.94 μmol/g and the cherry tomato (SM131) showed a moderate level of 21.02 μmol/g (Table 2). On average, 10-week-old *S. galapagense* accessions had 45% greater acylsugar concentrations than 8-week-old plants.

The leaflet area varied significantly (*P*<0.001) among *S. galapagense* accessions, ranging from 2.43 to 9.15 cm^2^. Leaflet area was significantly greater for cultivated tomato, with 14.36 cm^2^ than for all *S. galapagense* accessions. On average, cherry tomato leaflets were 10% larger than *S. galapagense* accessions (Table 2).

Correlations between trichome types and acylsugar concentration are presented in Table 3. Acylsugar concentration was positively associated with trichome type IV and negatively associated with trichome type V. The negative correlation between acylsugar and trichome type VI was only significant at the 10-week-old plant stage. No significant correlation was found between acylsugar and trichome type I. In addition, leaflet area was negatively correlated with trichome IV density of abaxial surface (*P*<0.001, Figure 2A) and acylsugar concentration (*P*=0.078, Figure 2B).

**Table 3.**
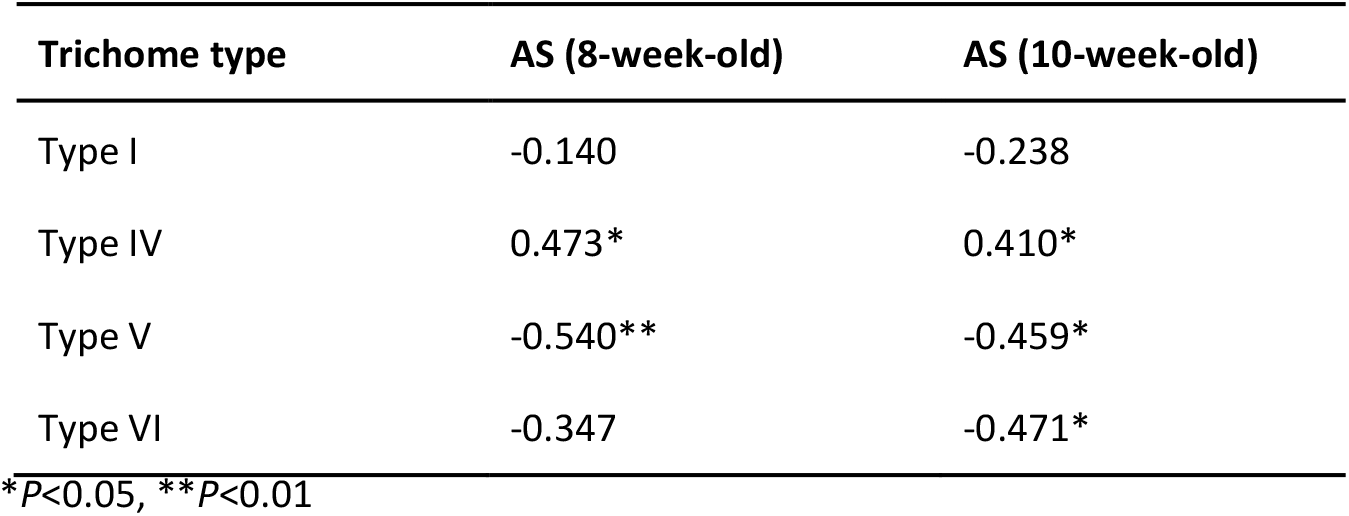
Correlations between trichome types and acylsugar (AS) concentration at 8-week (N=26) and 10-week (N=28) intervals on abaxial surface.

**Figure 2.**
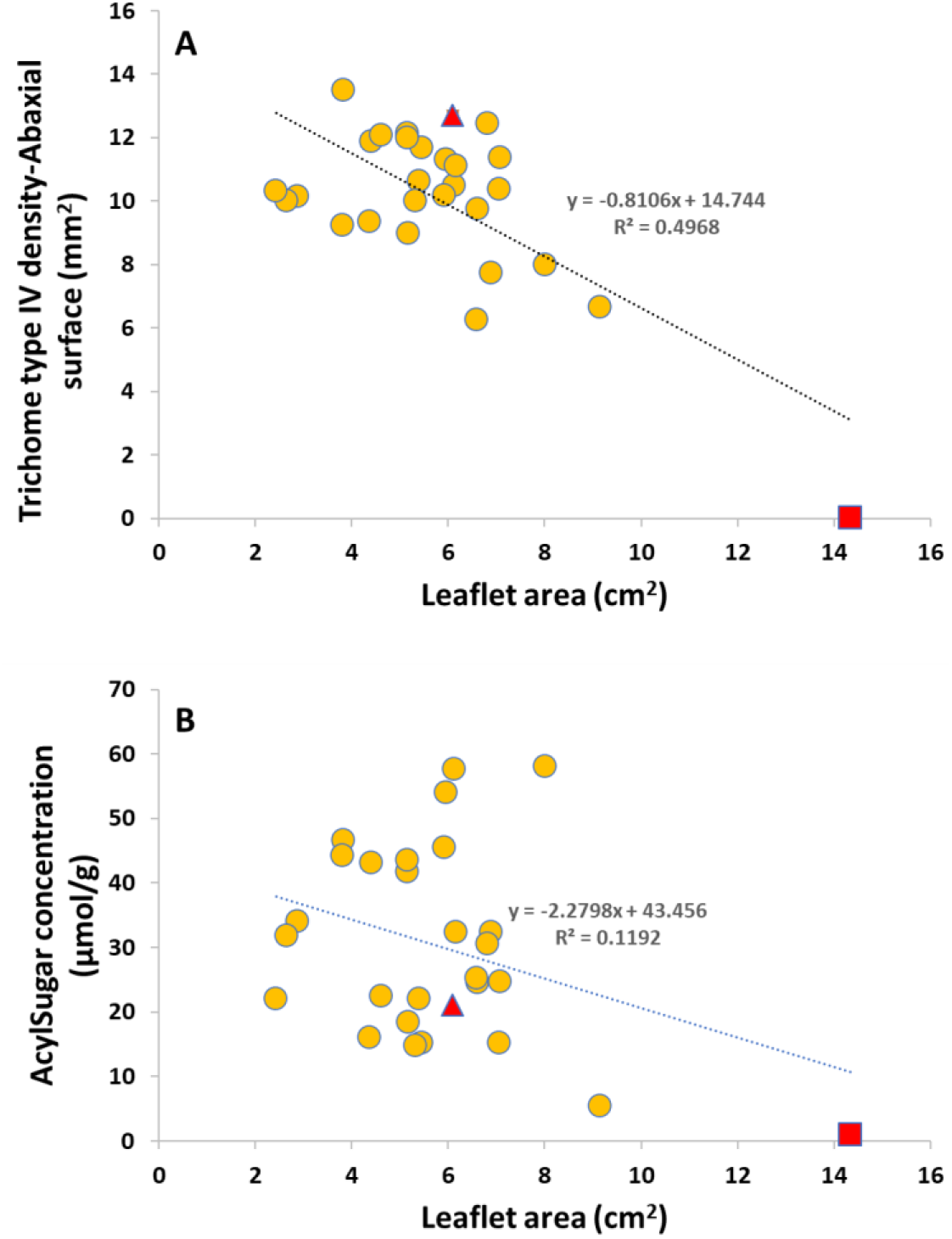
Relationships between trichome type IV (**A**) and acylsugar concentration (**B**) with leaflet area at 10-week-old seedlings (N=28). The red triangle and red square represent cherry and cultivated tomato cultivars, respectively.

The PCA plot (Figure3) showed three distinct clusters based on acylsugar, trichome type, and leaf area measurements. The first group consisted of *S. galapagense* accessions and the cherry tomato genotype (SM131) that was in correlation with high acylsugar concentration as well as high levels of trichome type IV on the adaxial surface. The cultivated tomato (CLN3682C) was separated from *S. galapagense* group mainly due to its greater leaflet area and high levels of trichome type V. One of the *S. galapagense* accessions, VI007099, was grouped closer to the cultivated tomato mainly due to its larger leaflet area compared to other accessions from the Galápagos Islands.

**Figure 3.**
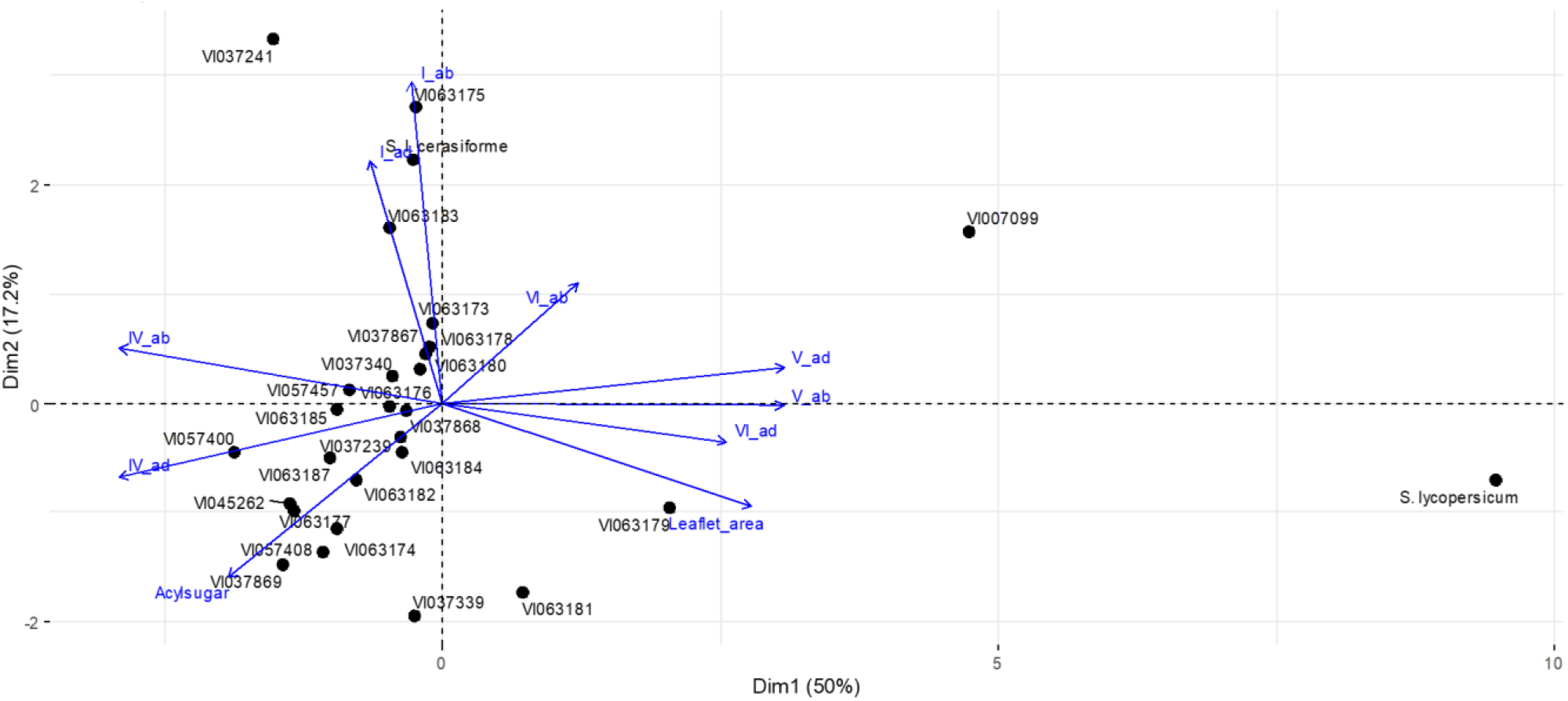
The biplot illustrates the principal components (PC) analysis for the 26 accessions of *S. galapagense* and two *S*. *lycopersicum* lines with measurements as vectors. Vectors that are close together are correlated in terms of the leaf measurements. ad, adaxial surface; ab, abaxial surface. I, IV, V, and VI refers to trichome types.

The DNA marker results are presented in Table S2. All accessions of *S. galapagense* showed both resistance bands at 46 and 40 bp for the *Wf-2* marker (indicated by “++”) while both *S. lycopersicum* cultivars (CLN3682C and SM131) showed the susceptible band at 87 bp (indicated by “-”). The *Wf-1* marker showed a resistance band at 139 bp (indicated by “+”). Both *S. lycopersicum* cultivars showed the susceptible band at 157 bp (indicated by “-”).

## Discussion

Here we screened a large germplasm collection of *S. galapagense* accessions revealing a considerable genetic variation in glandular trichome density and acylsugar concentration. Our results indicated that some accessions of this species are a promising source of high trichome types IV density and acylsugar concentration. We also found that tomato plants with smaller leaves had higher densities of trichome type IV and greater acylsugar concentration.

The selection of tomato plants with higher densities of glandular trichomes has shown to be an effective criterion to obtain superior resistance to insect pests. This has been widely proven in resistance to spider mites (*Tetranychus urticae*, Andrade *et al*., 2017; Rakha *et al*., 2017a; de Souza Marinke *et al*., 2022), whitefly (*Bemisia tabaci*,Rakha *et al*., 2017b; Lucatti *et al*., 2013), thrips (*Frankliniella occidentalis*,Escobar-Bravo *et al*., 2017), and cabbage looper caterpillar (*Trichoplusia ni*,Mymko and Avila-Sakar, 2019). These studies generally attribute the insect pest resistance to the excretion of acylsugars by the tip of glandular trichome type IV, thereby acting as a biopesticide.

Several studies have shown a large genetic diversity for trichome density and acylsugar concentration in tomato (e.g. Lucatti *et al*., 2013; Baier *et al*., 2015; Rakha *et al*., 2017b). These have allowed tomato breeding programs globally to exploit the available diversity and improve insect pest resistance cultivars. We screened a relatively larger number of *S. galapagense* accessions (26) compared to previous studies which characterised 10 accessions (Rakha *et al*., 2017b) or 11 accessions (Lucatti *et al*., 2013). This study seems to be the first to screen accessions VI037867, VI037869, VI045262, VI057457, VI063173, VI063178, VI063179, VI063181, VI063182, and VI063183 for insect pest resistance-related traits. Our study revealed a broader variation for trichome type IV, however, smaller values for acylsugar concentration compared to a similar study by Rakha *et al*. (2017b). The cherry tomato cultivar (SM131) was previously characterised by high trichome type IV density (unpublished data). Our results confirm that its trichome type IV density was higher than 96% of studied *S. galapagense* accessions (Table 2).

Not a strong correlation was observed between acylsugar concentration and trichome type IV at two different sampling times. Previous studies also reported a positive but moderate correlation between these two traits (Lucatti *et al*., 2013; Rakha *et al*., 2017b). A possible explanation for this could be the poor phenotyping of trichomes under the microscope, as counting the number of trichomes is an inherently delicate task. This difficulty highlights the need for high throughput methods to measure trichomes. Another explanation for the lack of strong correlation between trichome IV density and acylsugar could be that acylsugar production is not solely linked to trichome density but also their metabolic activity, whereby the same trichome types in different accessions, produce varying levels of acylsugar (Zhang *et al*., 2007; Bergau *et al*., 2015). Following this reasoning, isolated trichomes could be tested for metabolic activity through GC-MS as described for *L. hirsutum* by Fridman *et al*. (2005).

A negative correlation was observed between trichome type IV and leaflet area, a trend that has been reported in other plant species including *S. berthaultii* Hawkes (Pelletier, 2012) and silver birch (*Betula pendula* Roth) (Lihavainen *et al*., 2017). Using six cultivated tomato cultivars, however, Mymko and Avila-Sakar (2019) reported that the trichome density of expanded leaves did not differ significantly from that of unexpanded leaves at different growth stages. In our study leaflet area was the main driver to allocate tomato species into three different groups (Figure 3). The *S. galapagense* accession VI007099 representing leaf morphology between wild and cultivated tomato was clustered between two groups based on the trichome and acylsugar measurements. Our results suggest that ideal insect pest resistance tomato cultivars may have smaller leaflets.

The *S. galapagense* accessions originating from the Galápagos Islands have been exposed to dry and saline growing conditions (Pailles *et al*., 2020) and high insect pressure (Peck, 2008), thus representing a generous source of alleles that can be explored to improve biotic and abiotic stress. As this species can easily hybridize with cultivated tomatoes (Rick, 1961) it has been used as donors for stress tolerance genes, which could be transferred into commercial varieties by introgression breeding (Zamir, 2001). For example, VI037339 (LA1401) and VI007099 accessions have already been utilized as donors of high trichome IV density into modern cultivated tomato cultivars through interspecific crosses (Andrade *et al*., 2017; Rakha *et al*., 2017b; da Silva *et al*., 2019; Vendemiatti *et al*., 2021). However, in this study, these accessions were not among those with the highest trichome type IV density and acylsugar concentration. This could be due to the genotype-by-environment interactions as the experiments were carried out under different growing conditions.

From the analysis of DNA markers, we could see that most *S. galapagense* accessions were homozygous for *Wf-1* and *Wf-2* but neither *S. lycopersicum* cultivars. This suggests that the morpho-chemical measurements in this study were linked to the genetic background of *S. galapagense* accessions. However, the cherry tomato cultivar SM131 which showed high levels of trichome type IV and acylsugar did not amplify either band of the DNA markers. This was not surprising as those DNA markers were developed from an interspecific population derived from *S. galapagense* (Firdaus *et al*., 2013). We conclude that *Wf-1* and *Wf-2* may be more suitable to be used in genetic materials derived only from *S. galapagense*.

In conclusion, our study focused on screening a large *S. galapagense* germplasm, supporting breeding programs aiming to improve insect pest resistance in tomato using crop wild relatives. The ultimate goal is to develop tomato cultivars with insect pest resistance-related traits that help farmers to reduce pesticide use and produce a high-quality and chemical-free tomato crop.

## Acknowledgment

We would like to thank Yun-che Hsu (Grace) and Jean Lin for their kind assistance during the experiments. We also thank Dr. Roland Schafleitner (Flagship Leader, Vegetable Diversity & Improvement) and Dr. Maarten van Zonneveld (Genebank manager) for their valuable suggestions during the experiments. In addition, the first author would like to thank National Cheng Kung University (NCKU), Taiwan for its support.

